# Response of neuronal populations to phase-locked stimulation: model-based predictions and validation

**DOI:** 10.1101/2024.11.06.622295

**Authors:** Nima Mirkhani, Colin G. McNamara, Gaspard Oliviers, Andrew Sharott, Benoit Duchet, Rafal Bogacz

## Abstract

**Background:** Modulation of neuronal oscillations holds promise for the treatment of neurological disorders. Nonetheless, stimulating neuronal populations in a continuous open-loop manner can lead to side effects and suboptimal efficiency. Closed-loop strategies such as phase-locked stimulation aim to address these shortcomings by offering a more targeted modulation. While theories have been developed to understand the neural response to stimulation, their predictions have not been thoroughly tested using experimental data.

**Objective:** We aimed to test the predictions of a mathematical model regarding the response of neuronal populations to phase-locked stimulation.

**Methods:** Using a coupled oscillator model, we expanded on two key predictions describing the response to stimulation as a function of the phase and amplitude of ongoing neural activity. To investigate these predictions, we analyzed electrocorticogram (ECoG) recordings from a previously conducted study in Parkinsonian rats, and extracted the corresponding phase and response curves.

**Results:** We demonstrated that the amplitude response to stimulation is strongly correlated to the derivative of the phase response (*ρ* > 0.8) in all animals except one, thereby validating a key model prediction. The second prediction postulated that the stimulation becomes ineffective when the network synchrony is high, a trend that appeared missing in the data. Our analysis explained this discrepancy by showing that the neural populations in Parkinsonian rats did not reach the level of synchrony for which the theory would predict ineffective stimulation.

**Conclusions:** Our results highlight the potential of fine-tuning stimulation paradigms informed by mathematical models that consider both the ongoing phase and amplitude of the targeted neural oscillation.

## 1. Introduction

Application of brain stimulation techniques has gained momentum over the past few decades owing to their therapeutic potential [1,2]. Neural oscillations can act as anchor points in modulation of brain circuitry [3,4]. The association of particular network oscillations with different brain functions, as well as their implication in many neurological and psychiatric disorders, renders them suitable targets for stimulation [5–8]. Successful manipulation of neural oscillations for the desired outcome requires clear answers to where, how, and when to stimulate [9–11]. The first question has been extensively researched to identify the target site, which varies significantly based on the engaged networks [12–16]. To address how and when stimulation should be applied, a variety of closed-loop strategies have been proposed, where features of the ongoing oscillation serve as feedback for the stimulation [17–19].

Among closed-loop techniques, phase-locked stimulation has shown promise in achieving a controlled modulation [19–22]. In this approach, stimulation pulses are triggered at certain phases of the ongoing oscillatory activity. Neuromodulation and plasticity effects obtained through precise timing of the pulses have been shown to be bidirectional, enabling amplification versus suppression of the oscillations [23,24] and synaptic potentiation versus depression based on the target phase [25,26]. This feature not only results in higher control and in turn more efficient stimulation policies but may also explain the heterogeneity observed in many open-loop stimulation paradigms. Additionally, due to interactions between different brain rhythms through mechanisms such as phase amplitude coupling, phase-based modulation of an activity can bring about cross frequency changes [27,28].

Despite growing interest in phase-locked stimulation, two main bottlenecks of completely different natures have hindered further application of this strategy. Firstly, from a technical standpoint, real-time tracking of signal properties at the resolution of milliseconds is challenging. Thanks to recent technological advancements and developed algorithms, several studies have demonstrated the implementation of such fast brain-machine interactions for phase-locked stimulation in rodents [29,30], non-human primates [5,31], and humans [32,33]. Secondly, theoretical understanding of how the state of a network oscillation at the stimulation time, i.e. its phase and amplitude, modulates the response remains incomplete, often resulting in an extensive search during stimulation sessions to achieve the desired effect. There have been several theoretical studies proposing optimal closed-loop stimulation policies [34–37]. However, the predictions made by these studies have not been thoroughly validated with experimental data, which severely limits their applicability.

Mathematical models based on coupled oscillators are suitable candidates for bridging this gap due to their ability to replicate neural oscillations [34,38–40]. The Kuramoto model of coupled oscillators, in particular, offers a great advantage for studying phase-locked stimulation by adopting a phase-based description of neural oscillators [41,42]. As a result, network dynamics can be explicitly modeled as a function of individual oscillator’s phases, which evolve over time based on their interactions with each other and their corresponding natural frequencies. This relatively simple yet insightful model expresses the collective behavior of oscillators in terms of a mean phase and network synchrony, directly proportional to the amplitude of oscillatory activity [35]. Since the response to stimulation is shown to depend on the phase [20,29] and amplitude [35,43], predicting the network response to stimulation as a function of these two quantities may potentially provide clinically translatable predictive power, especially for stimulation-based treatments received by patients suffering from Parkinson’s disease (PD) or essential tremor (ET) [44,45]. The gained insight could also pave the way for combining phase-locked stimulation with adaptive stimulation—another closed-loop strategy that adjusts stimulation based on the amplitude of ongoing activity [46,47]—merging the best of both approaches.

Here, we aim to expand on the predictions introduced in [35] and test them using previously collected experimental data from [29]. We start by outlining the applied methodology and then introduce the predictions derived from the reduced (mean-field) Kuramoto model regarding the role of ongoing oscillations’ phase and amplitude in response to stimulation. Each theoretical prediction is then tested separately against the electrocorticogram (ECoG) measurements of Parkinsonian rats subjected to phase-locked stimulation at the Globus Pallidus (GPe). The original Kuramoto model is deployed where the reduced model falls short in accounting for different effects in the response. Using this analysis approach, we first demonstrate a strong correlation between the amplitude response curve (ARC) and the derivative of the phase response curve (PRC), where the ARC and PRC represent changes in amplitude and phase as functions of the stimulation phase, respectively. We then elaborate on the role of amplitude in magnitude of the stimulation effect, showing that the largest effects can be attained by stimulation at intermediate values of network synchrony. Below this peak, where most brain networks operate, the response is characterized by a slight drop and relatively stronger amplification compared to suppression. Taken together, these findings bridge the gap between theory and experiments, unveiling an opportunity to manipulate neural activities in the desired direction more reliably.

## 2. Materials and methods

To investigate the effects of stimulation, we introduce our modeling approach and describe the previously collected dataset used to validate the model’s predictions. We also detail the techniques employed to link theory and experiments.

### 2.1. Modeling framework

The Kuramoto model of coupled oscillators was used to model the oscillations arising from the activity of a neuronal population. In this framework, the network dynamics are described through phases that reflect self-sustained oscillations of weakly coupled oscillators [48]. We employed such a network model to analyze how external stimulation affects the network activity of a population. In this context, neurons or neural microcircuits with periodic behavior can be regarded as oscillators that interact with each other [35], collectively giving rise to the network activity often recorded in experiments as local field potentials (LFPs) or ECoG oscillations (Fig. 1A,B) [49,50].

**Fig. 1.**
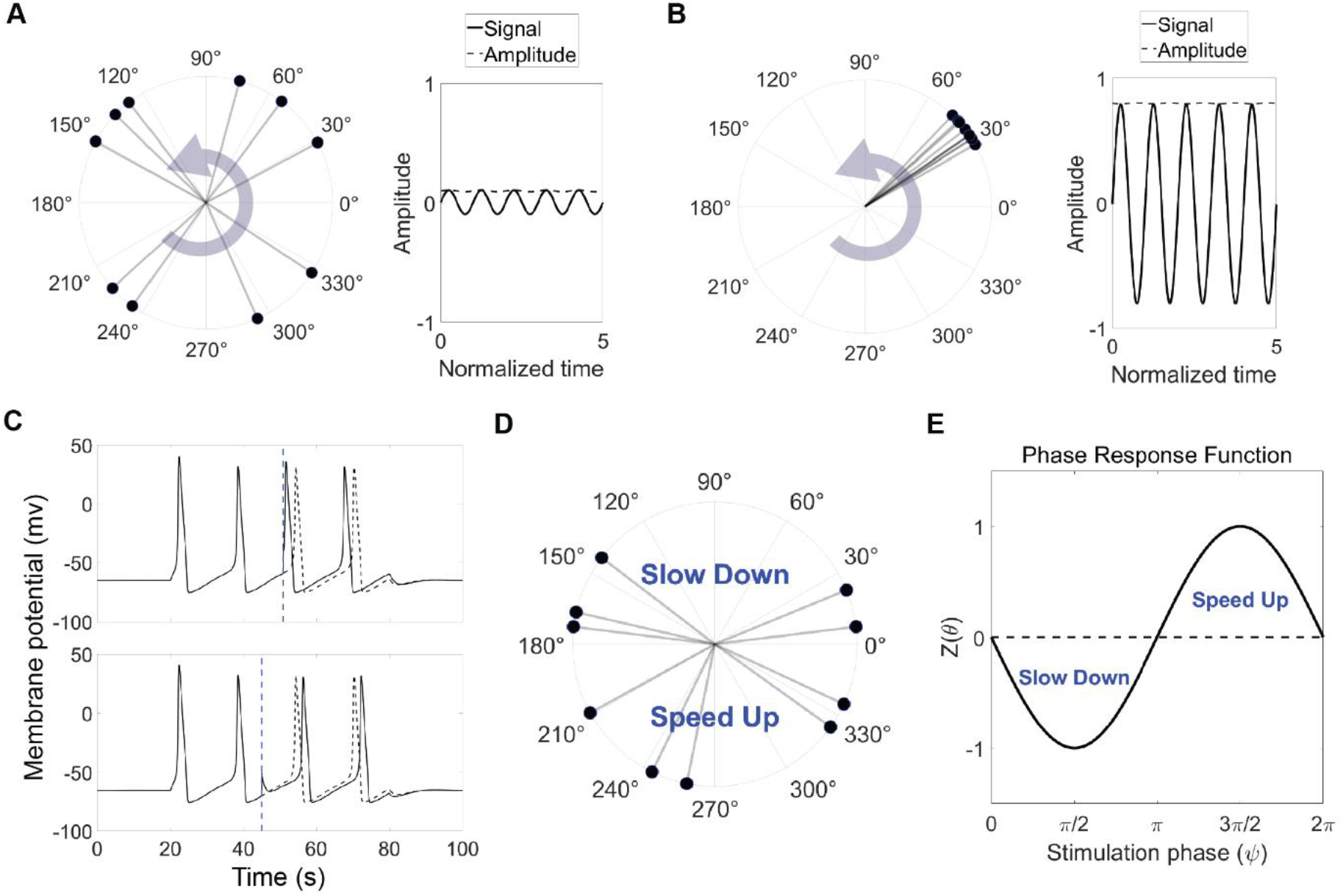
The Kuramoto model of coupled oscillators to model oscillatory neural activity. **A, B**. Snapshots of two example sets of coupled oscillators with low (A) and high (B) synchrony levels. Dispersed oscillators result in small oscillatory signal and therefore low amplitude, while packed oscillators represent large oscillation amplitude. **C**. Shift of spiking in a Hodgkin Huxley model as a result of external stimulation. Depending on the stimulation time with respect to the spiking cycle it can lead to phase advance (top) or delay (bottom). The dashed lines represent the spiking behavior in the absence of stimulation. **D**. Schematics of the biphasic response behaviour incorporated in the model. **E**. Phase response function of an individual oscillator (here, −*sinθ*).

To assess the impact of external stimulation on these networks, one must make an assumption about how individual oscillators respond to stimulation. Neurons may vary in their phase response depending on their type and various regulating factors [51–53]. We adopted the classic case compatible with the Hodgkin Huxley model, where a spiking neuron exhibits a biphasic response featuring both phase delay and advance regions [51]. This response behavior, known as type II, has been observed experimentally [54,55], and characterized by a slow-down region after the spiking during the refractory period and a speed-up region closely before the spiking [56] (Fig. 1C). We used *Z*(*θ*) = −*sinθ* as a simple phase response curve that satisfies these conditions (Fig. 1D,E).

### 2.2. Full Kuramoto model

Dynamics of a finite number of coupled oscillators with noise are governed by [57]:

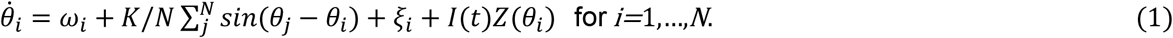

This set of differential equations describes how the phase of each oscillator, *θ*_*i*_, evolves in time while interacting with other oscillators through a global coupling constant *K* and subject to external stimulation *I*(*t*) and independent white noise *ξ*_*i*_:

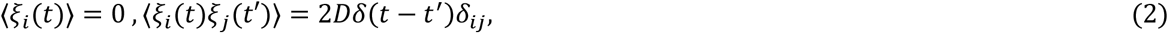

where *D* represents noise intensity. *δ* and *δ*_*ij*_ are the delta Dirac and Kronecker delta functions, respectively.

To investigate macroscopic properties of these networks, an order parameter is defined as follows:

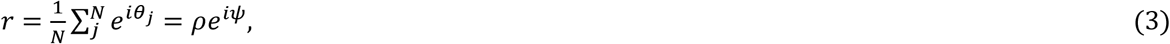

which describes the network activity in terms of the level of synchrony, ρ (ranging from 0 to 1), and the mean phase, ψ. It can be shown that the experimentally measured oscillation amplitude is proportional to the value of synchrony [35] (further details available in the Supplementary Section 4).

Numerical simulations were conducted in MATLAB using the Euler method with a time step *dt* = 0.0005 to discretize the system in time. The natural frequencies of oscillators were randomly sampled from a Cauchy distribution with a mean frequency of *ω*_0_ and a width of *γ*. To simulate phase-locked stimulation, conditions similar to those in [29] were applied. In each stimulation block, a target phase was chosen, and a pulse was delivered when the calculated mean phase crossed this target and more than 80 % of a beta cycle had elapsed since the previous pulse.

### 2.3. Reduced Kuramoto model

In the limit of an infinite number of oscillators and under certain assumptions regarding the distribution of natural frequencies, the collective behaviour of the network can be described in a simpler way solely by the time evolution of the order parameter [49,58]. The dynamics of the system are reduced to two differential equations governing the amplitude (synchrony *ρ*) and mean phase ψ of the network:

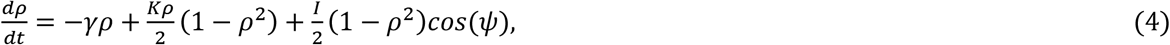

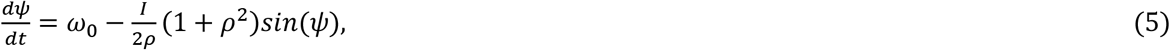

where *γ* represents the width of the natural frequency distribution that is centered around *ω*_0_. The last terms in the amplitude and phase equations represent the instantaneous population ARC and PRC, respectively.

In the absence of noise, such networks would reach a steady state condition with fixed values of *ρ*. However, in real networks, the oscillation amplitude fluctuates due to finite size effects, noise, and changes in coupling resulting from synaptic plasticity. Nonetheless, the reduced model can be seen as a phenomenological platform that provides intuitions and preliminary predictions. Accordingly, we employed this model as the basis for generating predictions, which were then further refined using the full model to partially capture the missing effects in a reduced model.

### 2.4. ECoG recording from Parkinsonian rats

In order to test the validity of the theoretical predictions, we used ECoG recordings collected from rats in [29]. In this study, in brief, rat models of PD were created through unilateral lesions of the dopaminergic neurons in substantia nigra, resulting in pathologically elevated beta activity in the cortico-basal ganglia network. Stimulating electrodes were then implanted in the globus pallidus (GPe), and activity was recorded using electrocorticogram (ECoG). Using a real time implementation of the phase tracking algorithm “Oscilltrack” [59], each subject underwent phase-locked stimulation at eight equally-spaced target phases based on the ongoing beta signal. Each trial was targeted at a specific phase and consisted of 10-14 stimulation blocks, each lasting 20 sec and separated by 5-sec off-epochs where no stimulation was applied (Fig. 2A). Full details are presented in [29].

**Fig. 2.**
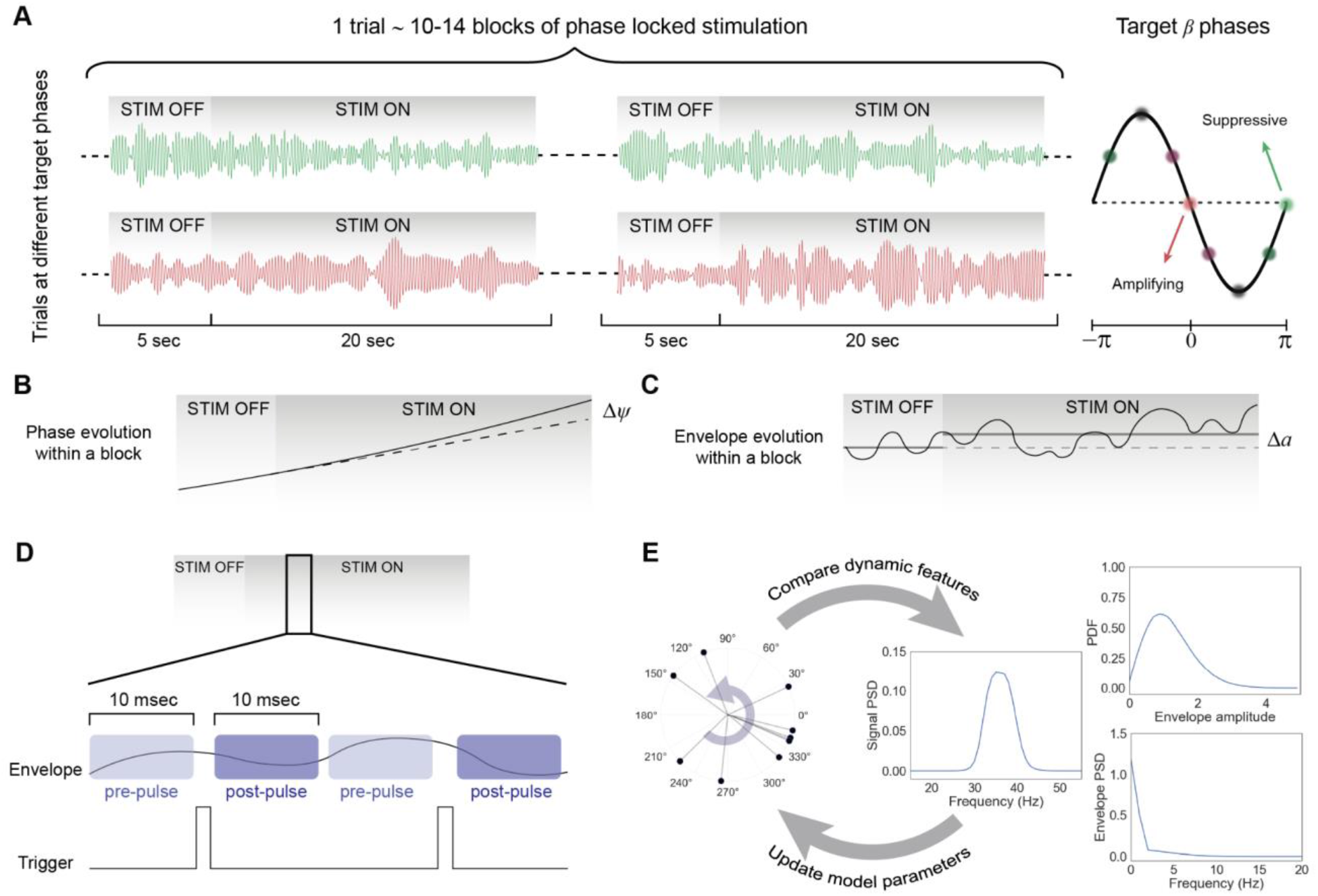
Analysis of experimental data for testing theoretical predictions. **A**. Summary of phase-locked experimental trials conducted in Parkinsonian rats in [29]. Each trial targeted a certain phase and was divided into 10-14 blocks. Two example signals, one representing stimulation at a suppressing phase (green) and one at an amplifying phase (red) are shown. **B**. Schematic of block-based quantification of the phase response **C**. Schematic of block-based quantification of the amplitude response. **D**. Schematic of pulse-based quantification of the amplitude response. **E**. Model fitting algorithm flowchart. Dynamic features of the subject-specific experimental signal were fed into an optimization solver which updates model parameters to minimize the difference between the model and experimental signals.

ECoG recordings were obtained at a sampling rate of 20 kHz. Stimulation artifacts were initially removed by interpolating the signal from the start of the electrical impulse to 1.5 ms after. The resulting signal was then downsampled to 2 kHz using an anti-aliasing filter. A 4^th^ order bandpass Butterworth filter was subsequently applied to the downsampled signal. The Hilbert transform was then used on the filtered signal to extract the envelope amplitude and phase of the beta oscillations.

### 2.5. Experimental response curves

The primary approach to extract the experimental ARC and PRC was the block-based method, in which the average behavior of the network during each 20-sec on-epoch was compared with the preceding 5-s off-epoch. More specifically, for the block-based ARC, the average Hilbert amplitude, ā, of the signal in each epoch was calculated, and the difference represented the amplitude change at the corresponding phase [20,60] (Fig. 2B). Evaluating this change for all target phases enabled us to reconstruct the experimental (block-based) ARC for each animal:

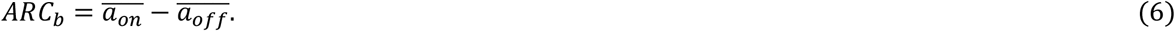

To calculate the block-based PRC, phase trajectory in the 5-sec off-epoch was used to fit a linear model for the phase evolution. Using this model, the expected phase of the system under no stimulation at the end of the 20-sec epoch, 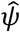, could be estimated. The difference between this estimated phase in the absence of stimulation and the actual phase, ψ, as a result of the stimulation was then normalized by the number of pulses in the on-epoch (Fig. 2C). This normalized change was calculated for all target phases, similar to the ARC, to establish the (block-based) PRC for each animal:

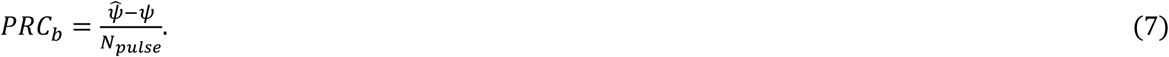

Considering the different sites for stimulation (GPe) and recording (cortex), as well as variability across animals, all curves were phase-aligned based on the most suppressive phase for the purpose of group analysis.

When quantifying the size of amplitude change under different oscillation amplitudes, the method above averages the oscillation over a relatively long period compared to the beta cycle’s time scale. To capture more transient changes in the amplitude, we also employed a pulse-based approach. In this technique, the average amplitude within 10 ms before and after each pulse was used to establish the amplitude response as a function of the pre-pulse amplitude (Fig. 2D).

A custom MATLAB script was developed to process the experimental recordings and extract the response curves. Statistical tests and additional data visualizations were carried out using Python-based packages. Pearson’s correlation coefficients (*R*) were calculated to assess the relationship between the ARC and the PRC derivative. Statistical significance of phase dependence in individual ARCs and PRCs was examined using one-way analysis of variance (ANOVA). The relationship between the correlation strength *R* and the resulting *p*-value from ANOVA was also quantified using the Spearman correlation coefficient (*r*_*s*_).

### 2.6. Model fitting

To test the prediction based on oscillation amplitude, it was necessary to estimate the network parameters that could reproduce relevant features of the ECoG recordings used for this study. An optimization-based model fitting algorithm was developed in MATLAB to fit the finite Kuramoto model to individual subjects. The algorithm received three dynamic features of the signal [60]: power spectrum density (PSD) of the signal, probability density function (PDF) of the envelope amplitude, and PSD of the envelope amplitude (Fig. 2E). These features embed the statistics of the signal intensity along with the temporal variations of both the signal and its amplitude [61]. It then employed MATLAB’s surrogate optimizer (surrogateopt) with batch update interval 1 to minimize the following error:

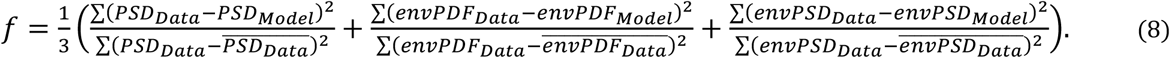

Given the different scale of the measured values and the model network activity, both experimental and simulated signals were z-scored to ensure comparability. PSDs were calculated using Welch’s method with frequency resolution of 1 Hz (1-sec window length) and 50 *%* overlap. The optimization output provided values for four network parameters: mean frequency *ω*_0_, width of the distribution *γ*, coupling constant *K*, and standard deviation of the noise σ. The maximum number of function evaluations for the surrogate optimization was set to 500. Network simulations at each optimization step were carried out with *N* = 200 oscillators which were randomly sampled from a Cauchy distribution with the mean *ω*_0_ and width *γ*. Each set of parameters was simulated 10 times to account for different realizations of noise, and the dynamic features from the resulting signals were averaged to calculate the optimization error.

A parameter recovery study was also performed using synthetic data to investigate whether the values obtained from the fitting procedure for individual parameters are separately identifiable with respect to network behavior. This series of simulations and optimizations were performed with fewer oscillators (*N* = 50) and lower frequency resolution for PSDs to reduce computational cost while still capturing the model’s generalizable features.

## 3. Results

The state of simple oscillatory systems can be summarized by their phase and amplitude (Fig. 3A). Hence, a clinically relevant predictive power may arise from studying the response to stimulation as a function of these two quantities tracked from signals of interest (e.g. tremor in ET or beta in PD). We first introduce the predictions made by the reduced model regarding the phase and amplitude dependence of the response to stimulation. We then examine the correlation between the PRC and ARC in the data from Parkinsonian rats. Finally, we compare amplitude-dependence in the experimental response with simulations of the best-fitting Kuramoto models.

**Fig. 3.**
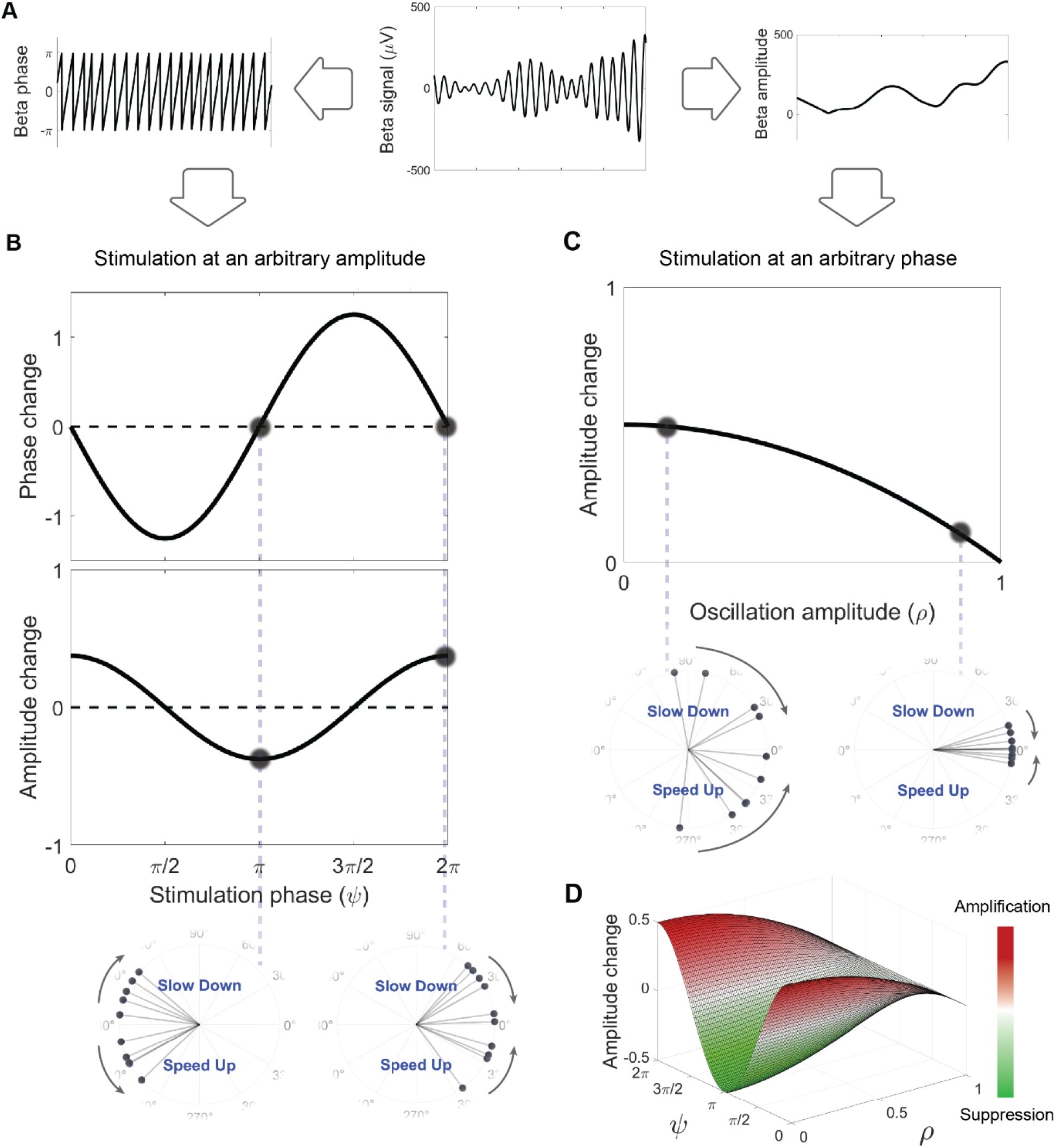
Theoretical predictions of the network response by the Kuramoto model. **A**. Phase and amplitude as the main signal properties. Predictions are based on the network’s state in terms of these properties. **B**. Model-based phase dependence of the network response. ARC follows the derivative of PRC. Two oscillators’ snapshots depict extreme cases of maximum suppression (left) and maximum amplification (right). **C**. Model-based amplitude dependence of the absolute change. Two oscillators’ snapshots illustrate stimulation-induced changes in networks with high (left) and low (right) synchrony. **D**. Combined effect of signal’s phase and amplitude on the amplitude response.

### 3.1. Introducing predictions from the reduced model

First, we focused on the phase dependence of the response behavior (Fig. 3B). Given a specific phase response function for individual oscillators, the reduced model predicted that the population PRC which represents the phase response of the network mirrors the form of the individual oscillators’ response function [35]. More importantly, the amplitude response of the population summarized by the ARC will be correlated with the derivative of the PRC. To develop an intuition about this prediction, two extreme scenarios of maximum suppression and maximum amplification can be helpful. In the former case, when stimulating the network at the mean phase of π, the trailing oscillators are in the “slow down” region of their cycle while the leading ones have entered the “speed up” regime (Fig. 3B, left inset). As a result, stimulation enlarges the gap between oscillators, causing a more desynchronized system. Conversely, in the maximum amplification scenario, stimulating at the mean phase of 0 causes leading and trailing oscillators to experience opposite effects, making them more tightly packed and thus synchronized (Fig. 3B, right inset). Hence, the maximum suppression and amplification correspond to phases where the absolute slope of the PRC is the largest.

Next, we sought to predict the network response in terms of amplitude dependence. In the reduced model, the last term of eq. (4) 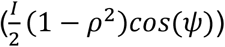 describes the instantaneous effect of a stimulation impulse, and the scaling factor of this term 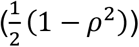 is plotted in Fig. 3C. This figure illustrates that the attainable absolute change in amplitude from stimulation drops continuously as a function of synchrony in the network. In other words, stimulation has the greatest effect when applied at low oscillation amplitudes and becomes less effective at large amplitudes. An intuition for this prediction can be obtained by looking at two ends of the synchrony spectrum. Any change in network synchrony requires differential effects of stimulation on the oscillators which leads to an increased or decreased gap between them. In a network with low synchrony, the high dispersion among oscillators allows for the maximum attainable change as a result of stimulation (Fig. 3C, left inset), whereas in a highly synchronized system, all oscillators experience nearly the same change, leading to minimal impact on the collective synchrony (Fig. 3C, right inset).

Examining eq. (4) for amplitude in the reduced model, the last term represents the combined effect of phase and amplitude which can be visualized as a 3D surface (Fig. 3D). However, for a more detailed assessment of the predictions and better understanding of the model’s strengths and witnesses, we tested each component separately using the experimental data.

### 3.2 Correlation between the PRC derivative and ARC

To test the theoretical predictions regarding ARC and PRC, we extracted the corresponding curves from the animal data. Analysis of changes in beta power as a function of target phase in [29], revealed nearly antiphase maximum amplification and suppression in all animals. This general trend suggests that *Z*(*θ*) = −*sinθ* can be viewed as a reasonable assumption for phase response function of individual oscillators (see eq. (1)). Nevertheless, ARC and PRC curves for each animal enabled a more comprehensive analysis of the predictions. The block-based method, described above, was employed to calculate the phase and amplitude changes in the high-beta activity as a function of the phase of stimulation. The average PRC, pooled across all animals, exhibited the previously described “slow down” and “speed up” regions for the population activity (Fig. 4A). The corresponding ARC also confirmed antiphase maximum suppression and amplification with smooth transitions in between (Fig. 4B). More importantly, the core prediction of the model, which posits a correlation between ARC and PRC derivative, was examined by establishing the derivative curve calculated through central differencing (Fig. 4B). Comparing ARC and PRC derivative revealed a negative correlation, in agreement with the model’s prediction.

**Fig. 4.**
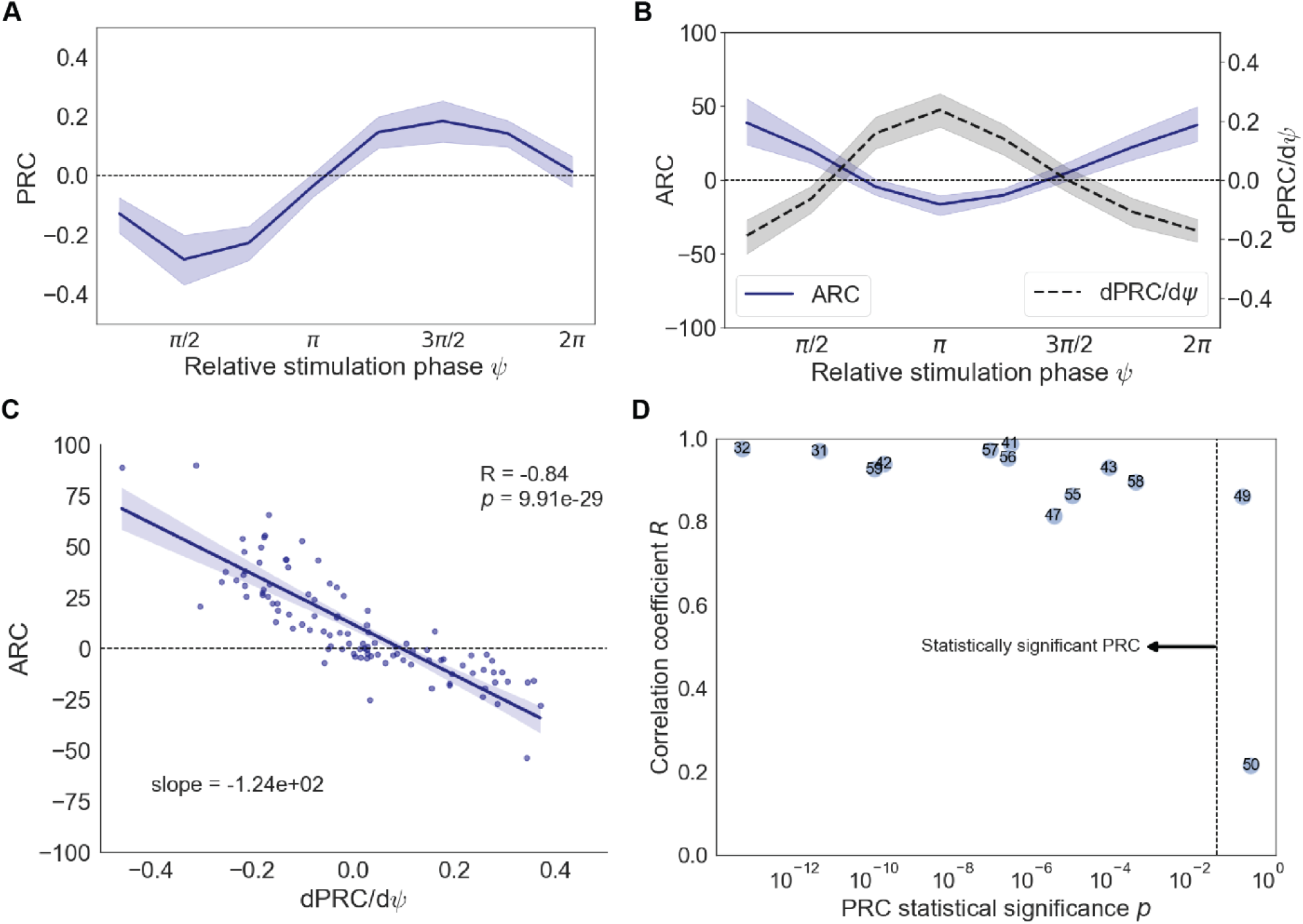
Experimentally validated relationship between ARC and PRC. **A**. Average PRC pooled across all animals. The average response exhibits phase advance and delay for the population activity. **B**. Average ARC (solid line) and PRC derivative (dashed line) pooled across all animals. The average amplitude response in rats is negatively correlated with the derivative of the phase response. **C**. ARC as a function of PRC derivative for 13 animals and 8 target phases. **D**. Relationship between the correlation coefficient and the significance of phase dependence within animals. Subjects with more statistically significant phase dependence tend to have a stronger correlation between ARC and PRC derivative.

To further quantify this correlation and its variability across animals, we examined the phase and amplitude responses for 13 individual animals at each of the 8 target phases (individual response curves available in Fig. S1). The data across all animals and phases showed a tight distribution around a line with a negative slope, resulting in a high correlation coefficient (*R* = 0.84) which underscored the validity of the predicted relationship (Fig. 4C, individual correlations available in Fig. S2). To assess how reliably the amplitude response can be predicted given a specific PRC, the relationship between the correlation strength, *R*, and the presence of an effect of phase in the PRC was explored. The latter was represented by the *p*-value from the statistical tests where lower values indicate significant phase dependence in the PRC. Plotting these values for different subjects revealed an interesting trend regarding variability across animals (Fig. 4D). All subjects with statistically significant PRCs exhibited a strong correlation with their amplitude response (*R* > 0.8). Notably, the phase response of the only subject lacking this correlation did not reach the significance threshold. Additionally, subjects with a higher effect of phase tended to show stronger correlations (*r*_*s*_ = −0.71, *p* = 6.7*e* − *3*). These results suggest that when certain model assumptions are met, —specifically, when phase dependence is present in the response— a tight correlation between ARC and PRC derivative may yield clinical insights when evaluating stimulation outcomes.

### 3.3. Contributing factors in amplitude modulation of the response

Following the study of phase dependence, we proceeded with analyzing how the network response is influenced by the ongoing oscillation amplitude. The previously described prediction on the dependence of stimulation effects on the ongoing amplitude (Fig 3C), was derived from the instantaneous effect of stimulation, i.e. the effect was defined as difference between amplitude of oscillations just after and just before the pulse. However, to understand longer term effects, one needs to also consider the dynamics of the system between the pulses. The changes in the oscillations amplitude in the model are described by eq. (4). It states that the amplitude ρ is not only influenced by the stimulation term but also depends on the coupling *K* and the distribution width *γ*, which together determine how amplitude evolves in subsequent time steps. In other words, it is not possible to study the longer-term response as a function of amplitude without considering the intrinsic network parameters. In addition, oscillation amplitude is naturally bounded by the minimum and maximum levels of synchrony (0 < ρ < 1).

To develop an intuition about the interaction of these contributing forces in amplitude modulation, a seesaw analogy can be useful (Fig. 5A). Each position of a seesaw corresponds to a specific balance between two opposing forces: one pushing the system towards synchrony and the other causing desynchronization. Within the Kuramoto framework, intrinsic noise *D* and width *γ* of the natural frequency distribution, and external stimulation at phases around the mid-ascending part of the cycle tend to reduce the synchrony of the network, tipping the balance towards lower ρ values (Fig. 5A, top). On the contrary, coupling *K* in the system and stimulation at phases around the mid-descending part shift the balance in favor of higher ρ values by enhancing the network synchrony (Fig. 5A, top). Furthermore, analogous to a real seesaw that is constrained at both ends, there are lower and upper bounds on how the force imbalance is reflected in the network (Fig. 5A, bottom).

**Fig. 5.**
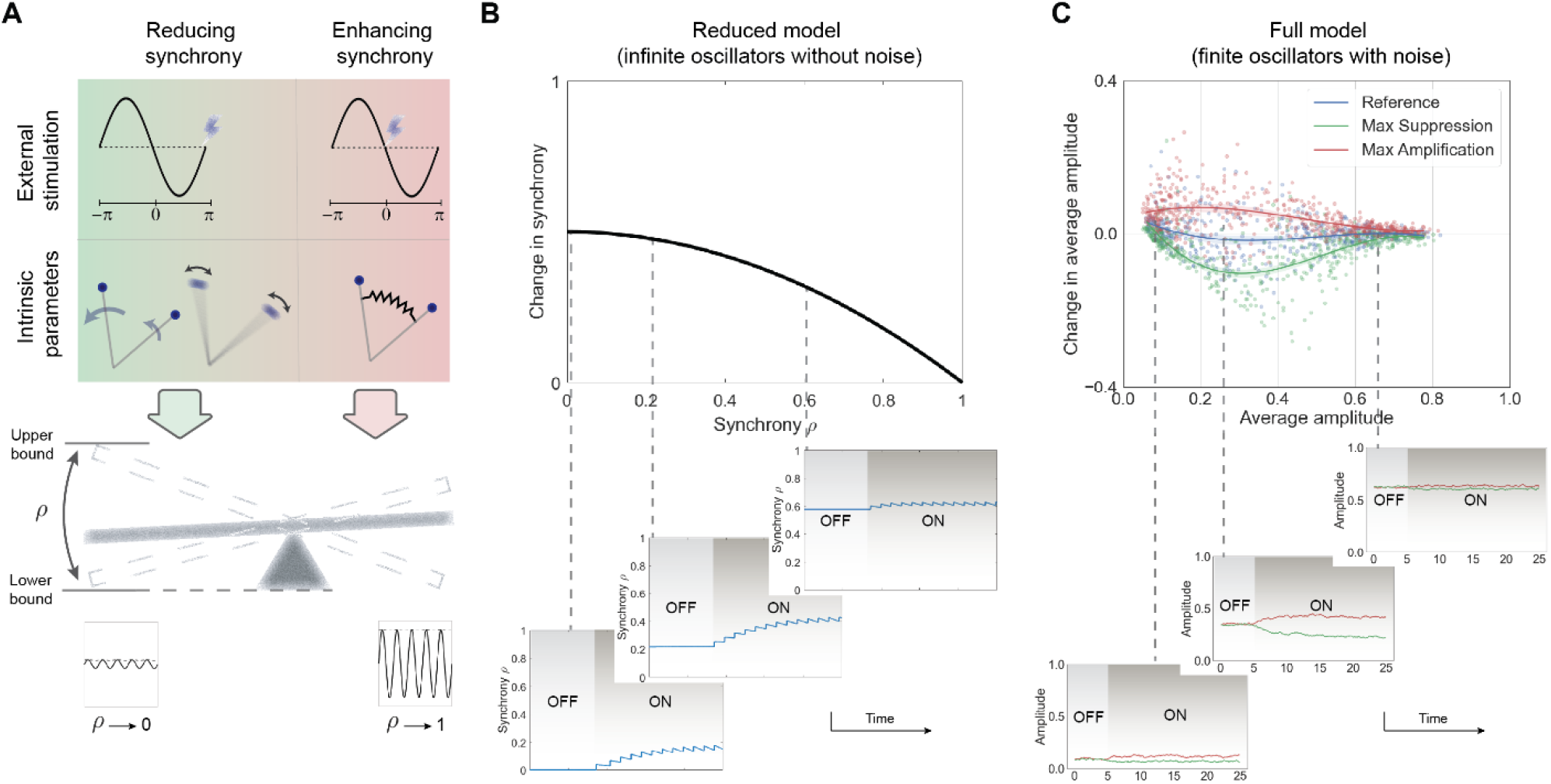
Contributing factors in amplitude modulation of the network response. **A**. A seesaw analogy of a network with two opposing forces and natural upper and lower bounds. External stimulation interacts with intrinsic parameters to determine the synchrony level in a constrained system. **B**. Stimulation-induced instantaneous disturbance as a function of oscillation amplitude. Three examples of synchrony evolution in the reduced model, each representing different steady-state synchrony levels. The extent of change in network synchrony as a result of stimulation depends on the size of the induced disturbance as well as network tendencies at that specific amplitude. **C**. Simulated block-based change in oscillation amplitude as a function of amplitude prior to stimulation in the full model. Three instances of amplitude evolution within a block illustrate the combined effect of stimulation-induced disturbance, intrinsic network tendencies, and synchrony boundaries.

As mentioned earlier, the introduced Kuramoto model’s prediction for absolute change as a function of oscillation amplitude highlights only the stimulation-dependent term in the time evolution of amplitude (Fig. 5B, top), without taking into account network dynamics influenced by intrinsic parameters. The reduced model offers an initial insight into the interaction of these contributing factors. In asynchronous networks (Fig. 5B, left inset), relatively high stimulation-induced perturbations are partially offset by network’s tendency to return to its steady-state with a low synchrony during the intervals between the pulses. As the network begins transitioning to a partially synchronized state (Fig. 5B, middle inset), a slight decrease in the effect of individual stimulation pulses emerges, but the network is notably more susceptible to changes, reflected in a smaller decay between pulses, which translates into larger shifts in average network synchrony. Lastly, under substantial levels of synchrony (Fig. 5B, right inset), not only is the stimulation effect diminished, but the network again shows a strong tendency to maintain its steady state, leading to smaller net changes in synchrony. Such variations in network tendencies could be better understood by looking at its characteristic curve (Fig. S3).

To further characterize the absolute change as a function of amplitude and link the model’s prediction with the experimental data, evolution of oscillation amplitude within similar stimulation blocks was simulated (Fig. 5C, top). The size of impulse was adjusted to match the observed change in beta power in experiments. These simulations were performed using the finite model with stochastic oscillators, and curves corresponding to no stimulation, most suppressive, and most amplifying phases were generated. The combined effects of external stimulation and intrinsic parameters constrained by bounds on both ends were consistent with the above descriptions. The relative weight of each contributing factor at different synchrony levels can be better understood by examining the time evolution of amplitude across three distinct levels of synchrony (Fig. 5C, bottom). In a highly noisy/low coupling network, only the amplifying phase caused a small upward shift in amplitude, as the already low amplitude could not be significantly reduced by stimulation at the suppressing phase (Fig. 5C, left inset). Under intermediate levels of coupling, far from both bounds of synchrony, a two-sided stimulation effect emerged in the response (Fig. 5C, middle inset) which then disappeared under asymptotically high levels of coupling due to the small instantaneous effect of stimulation (Fig. 5C, right inset).

### 3.4. Amplitude dependence of the response in Parkinsonian rats

Having refined the prediction for the magnitude of the stimulation effect as a function of synchrony level, we aimed to test it by extracting the corresponding response behavior from the rat data. As outlined in the Methods section, the oscillation amplitude in the model represents a normalized amplitude corresponding to the level of synchrony. Therefore, to compare experimental curves with theoretical predictions, one needs to first estimate the network synchrony corresponding to the measured ECoG. The model fitting algorithm was employed to determine the subject-specific network parameters *ω*_0_, *γ, K, D*. A series of parameter recovery studies using synthetic data revealed that while individual parameters could not be reliably recovered (details available in the Supplementary Section 3), the resulting network synchrony derived from each set of parameters was recovered with reasonable confidence (Fig. 6A). As a result, rather than focusing on exact parameter values from the fitting output, the corresponding network synchrony has been reported which indeed is more directly relevant to the theoretical prediction. The fitting was performed to replicate three key dynamic features of the signal for each animal (Fig. 6B).

**Fig. 6.**
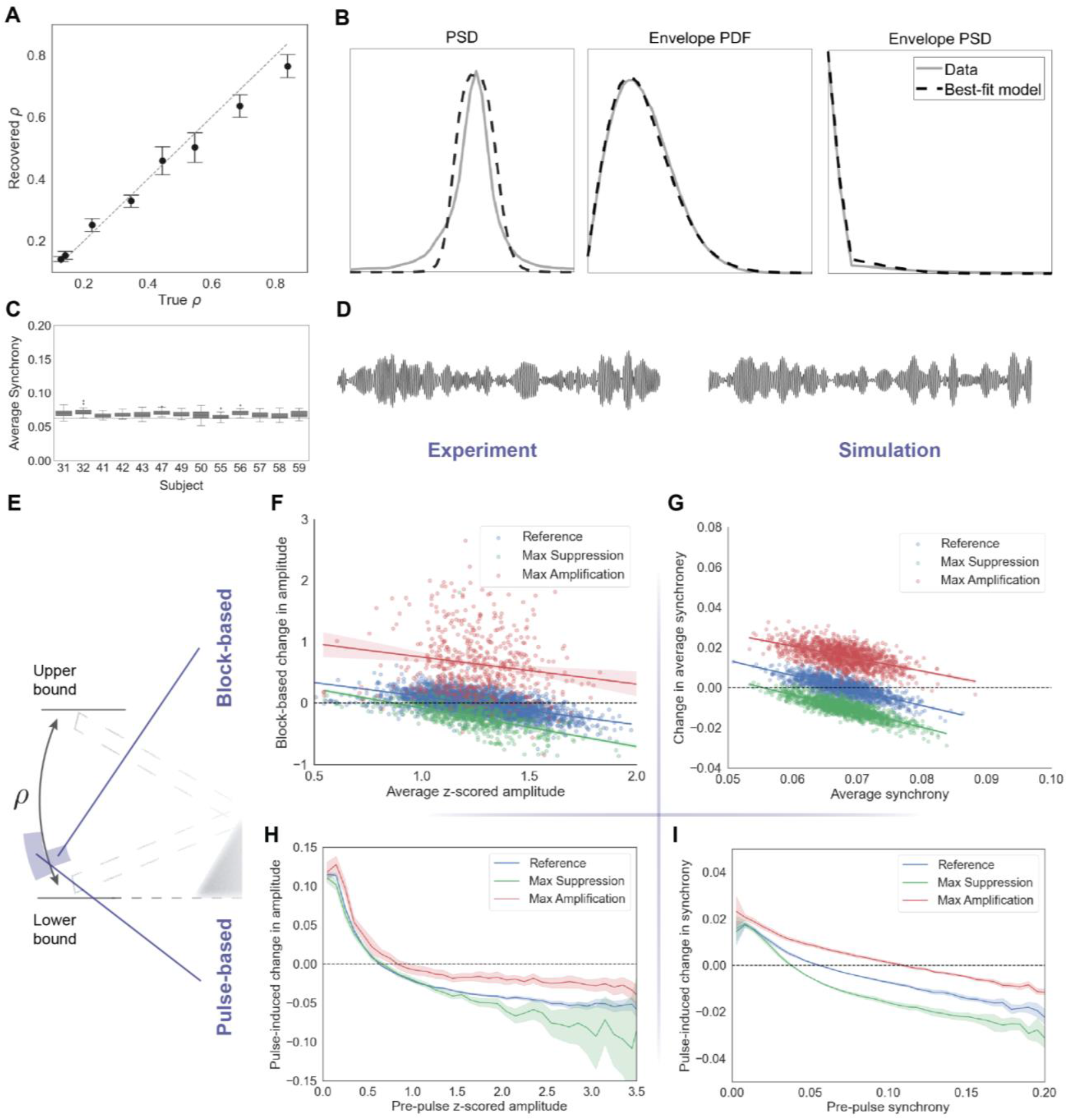
Network response dependence on oscillation amplitude. **A**. Parameter recovery using synthetic data. The model successfully recovers network synchrony from a set of simulated network activities. **B**. Results of the fitted dynamic features of the signal for an example animal. The power spectral density (PSD) of the signal and its amplitude, along with the probability density function (PDF) of the amplitude, were used to find the best-fitting parameters. **C**. Extracted mean synchrony for different subjects obtained by fitting the Kuramoto model to experimental recordings. The dashed line represents the mean synchrony when desynchronizing factors approach infinity. **D**. Example epochs of experimental recordings compared with simulated activity generated by the best-fit model. **E**. The relevant range of synchrony in the seesaw analogy. Block-based and pulse-based analyses provide different windows into the network synchrony. **F, G**. Block-based changes in amplitude as a function of pre-stimulation amplitude, demonstrating stronger amplification compared to suppression in this regime. **H, I**. Pulse-based changes in amplitude as a function of pre-pulse amplitude, revealing a drop in effect size at lower ends of the synchrony range through this quantification method.

Results of the model fitting implied that underlying networks producing the measured signals may possess very low sustained synchrony across all animals (Fig. 6C). To highlight that different combinations of network parameters can lead to the same oscillating behavior, reflected by the synchrony level, the fitting algorithm was run multiple times for each subject. Although output parameter values varied within each subject, all combinations consistently represented similar levels of network synchrony (Table S1). The resulting model networks produced beta oscillations similar to those observed in the experiments (Fig. 6D). Analytical estimates also confirmed that the data is consistent with low synchrony levels (details provided in the Supplementary Section 4).

Next, having identified the synchrony range of interest, we investigated the experimental data under two synchrony windows (block-based vs pulse-based variations) to assess whether the experimental responses aligned with the response of a model operating in that regime (Fig. 6E). As demonstrated in Section 3.3, low-synchrony networks are predicted to generally exhibit a higher propensity for amplification compared to suppression (Fig. 5C). By closely examining the blocks in the experiments, the amplitude change as a function of average amplitude prior to the stimulation epoch was obtained under three conditions: reference (no stimulation applied) plus the two phases achieving the highest amplification and highest suppression (Fig. 6F). At a given state, the inherent noise and finite number of oscillators caused a regression to the mean in the absence of stimulation. The model’s response, based on best-fit parameters, aligned with the experimental curves (Fig. 6G). The reference-subtracted regression lines reflected a significant increase in the stimulation effect during the suppressive phase, along with a more subtle, nonsignificant increase for amplification, and an overall more pronounced amplification in this regime (Fig. S7A).

Lastly, recognizing that using averaged amplitudes over on- and off-epochs narrows the analyzed synchrony window (higher and lower synchrony values are averaged out), similar amplitude dependence curves were also derived based on individual stimulation pulses instead of blocks. This approach accounts for a wider range of momentary synchrony levels that the network experiences. The experimental pulse-based curves generally exhibited similar trends to the block-based ones, except even smaller changes at the lower end and a peaked trend for the amplification (Fig. 6H). These differences were again in agreement with the curves extracted from the best-fit models (Fig. 6I). To visualize the net effects of stimulation, the corresponding reference line could be subtracted from the amplification and suppression lines (Fig. S7B).

## 4. Discussion

In this study, we elaborated on the predictions of a mathematical model based on coupled oscillators regarding the effects of phase-locked stimulation on a population activity. The model put forth predictions on how a neuronal population would respond to stimulation based on its current state in terms of phase and level of synchrony. We utilized a previously collected dataset from the study of phase-locked stimulation in rat models of PD to test those predictions.

For phase dependence, the prediction implied that the shape of ARC would follow the negative of the derivative of PRC, and all except one animal exhibited response behaviors consistent with the model’s prediction. This key relationship, validated for the first time in this study, has been investigated previously with different models. Using the Wilson-Cowan model, it has been demonstrated [60] that the phase shift between the ARC and PRC converges to π/2 in the linearised model. The phase shift was however larger than π/2 in the non-linear Wilson-Cowan model and in some of the data from patients with essential tremor. The predictive feature of the PRC derivative was also discussed in [62,63] through a noisy oscillators model, and in [36] with a network of conductance-based neurons. On the other hand, several studies [5,31,43] have adopted mathematical frameworks to explain the phase-dependent response to stimulation. Findings of these studies have been generally consistent with the explicit relationship between the ARC and PRC discussed here which is derived from a phenomenological model. In addition, when evaluating the effect of phase, it must be kept in mind that perfect phase tracking in practice is not feasible, especially at low amplitudes due to a lower signal-to-noise ratio. Consequently, quantifying the effect of phase based on the higher amplitude portions of the signal may provide a clearer perspective. Alternatively, collecting more data can help mitigate this issue by averaging out the variations caused by imperfect tracking, which was the case for the data used in this study. Overall, our results support the proposal that a prior estimate of the PRC (such as measurements conducted for cotrical [64], subthalamic [65], or pallidal neurons [51]) may be a useful tool for determining the suppressing or amplifying phase for closed-loop DBS without a full search of the parameter space [36]. This approach could provide valuable guidance for defining optimal stimulation parameters in clinical settings.

Amplitude dependence is a relatively unexplored aspect of the network response. Focusing solely on the instantaneous effects of stimulation, the theory suggests that stimulation should become ineffective at high network synchrony. However, the amplitude dependence was demonstrated to be more complex as other contributing factors such as network’s tendencies and bounds interact with stimulation-induced changes. These interactions would lead to a decay in the effect size of stimulation at both ends of the synchrony range. Moreover, distinct characteristic behaviors may emerge when amplifying oscillatory activity compared to its suppression, as exemplified by the stronger amplification observed in this study. This highlights the significance of determining the synchrony levels of the target network beforehand if the goal is to optimize stimulation efficiency based on ongoing amplitude. This could explain some of the observed differences in suppressing pathological activity in patients with PD compared to those with ET [19,66]. We proposed here that fitting the Kuramoto model to individual recordings could provide subject specific models, enabling tailored stimulation paradigms according to the subject and network under study. It is also important to note that the size of electrical impulses and the attainable modulation in clinical settings, which was simulated here to achieve comparable changes in power, could shift the location and intensity of peaks in the amplitude dependence of the response. The first-order trend, which is a general drop of the effect size with increasing amplitude, has been reported in several studies [43,67], and agrees with the intuition that the stronger the synchrony of a network, the harder it is to disrupt.

Beta oscillations in PD are considered an exemplar of pathological hypersynchrony. Therefore, it could be considered surprising that the stimulation effect did not drop at higher amplitudes in Parkinsonian rats. Importantly, however, high synchronization in the Kuramoto model represents almost complete alignment of individual oscillators (e.g. Fig. 3C, right). In the Parkinsonian brain, beta synchronization between neurons is massively elevated compared to healthy animals, where there is very little oscillatory synchronization [68–70]; However, if oscillators in the model represent individual neurons or ensembles of neurons in the basal ganglia circuit, these pathological levels of synchronization do not approach the levels of hypersynchronisation in the model. For example, in the subthalamic nucleus of Parkinsonian patients the maximum proportion of individual neurons that oscillate at beta frequency and/or are synchronized with cortical beta oscillations is around 60%, with a mean of 20-30% [71,72]. In the context of developing novel approaches for DBS in PD, this suggests that even highly patholophysiological levels of beta synchronization remain in the region where they remain responsive to modulation by phase-dependent stimulation. It remains to be seen whether this is also the case for pathophysiological activities with higher levels of synchronization, such as epilepsy.

With regards to limitations, although the model seems to capture the mean synchrony for subject-specific models that reproduce the ECoG recordings, it falls short of replicating the variability of oscillation amplitude observed in the animals, as seen when comparing the x-axes in Fig. 6F-I. This limitation could potentially be addressed by allowing for changes in the coupling as a result of synaptic plasticity, and/or using a more generalized coupled oscillators model where oscillators are allowed to vary in their amplitudes. Additionally, while the model also makes predictions about specific stimulation phases that lead to suppression or amplification given a specific response function *Z*(*θ*), only the correlation between ARC and the PRC derivative was tested due to the separate stimulation and recording sites in the experiments. Applying the developed framework on experimental data where sensing and stimulation has been conducted through the same electrode may facilitate further validation of the model’s prediction. Moreover, the focus here was placed on beta rhythms originated from basal ganglia which feature a bursty characteristic with very low sustained synchrony. Testing the model through other brain rhythms and networks will provide a more comprehensive image of the effects of phase-locked stimulation. Lastly, alternative methods of measuring the experimental ARC and PRC could lead to slightly different outcomes which is why analytical methods alone may not be sufficient to estimate network synchronies.

In summary, this study aimed to bridge the gap between theory and experiments by validating relatively straightforward yet powerful predictions. Such mechanistic understanding of the effects of stimulation could complement model free approaches like machine learning techniques to design more effective stimulation policies. The findings of this study highlight the significance of pinpointing the right time for stimulation, providing clinically translatable insights for optimizing closed-loop strategies.

## Data availability statement

No new data were generated in this study. The experimental data used here is available at http://dx.doi.org/10.5287/bodleian:9omadD7Pp. The developed codes for mathematical modeling and computational analysis would be made available on https://github.com/Bogacz-Group upon publication.

## Declaration of Competing Interest

C.G.M. and A.S. are inventors on a pending patent application related to the subject matter of this paper.

## Acknowledgments

The authors would like to acknowledge the use of the University of Oxford Advanced Research Computing (ARC) facility in carrying out this work. http://dx.doi.org/10.5281/zenodo.22558

## Funding information

N.M. and R.B. were supported by Medical Research Council Grant MC_UU_00003/1.

C.G.M. was supported by the Wellcome Trust (Sir Henry Wellcome Fellowship 209120/Z/17/Z).

A.S. and C.G.M. were supported by Medical Research Council Grant MC_UU_00003/6.

B.D. was jointly supported by the Royal Academy of Engineering and Rosetrees Trust under the Research Fellowship programme.

## CRediT authorship contribution statement

**Nima Mirkhani:** Writing – review & editing, Writing – original draft, Visualization, Validation, Software, Methodology, Investigation, Formal analysis, Conceptualization. **Colin G. McNamara:** Writing – review & editing, Investigation, Methodology, Funding acquisition. **Gaspard Oliviers:** Methodology. **Andrew Sharott:** Writing – review & editing, Supervision, Funding acquisition. **Benoit Duchet:** Writing – review and editing, Methodology, Supervision, Funding acquisition. **Rafal Bogacz:** Writing – review and editing, Supervision, Funding acquisition, Methodology, Conceptualization.

