## Supplementary material for "Response of neuronal populations to phase-locked stimulation: model-based predictions and validation"

### 1. Relationship between ARC and PRC in individual animals.

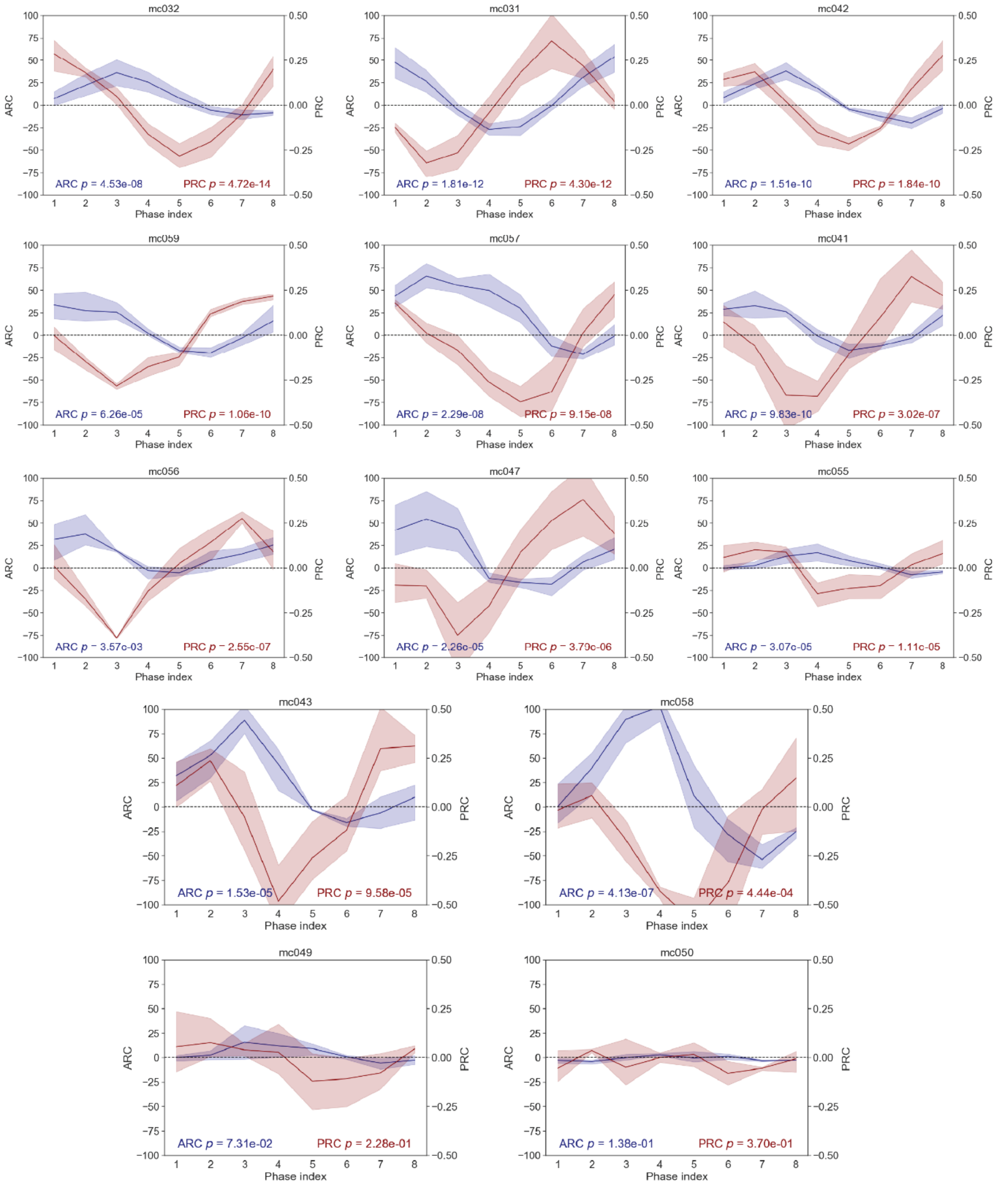


**Fig S1. Response curves for individual animals** (Blue: ARC, Red: PRC). For each curve, the corresponding p-values from the one-way ANOVA test are shown at the bottom. Plots are sorted according to the *p*-value of the PRC curves. Only two animals, mc049 and mc050, exhibit non-significant response curves.

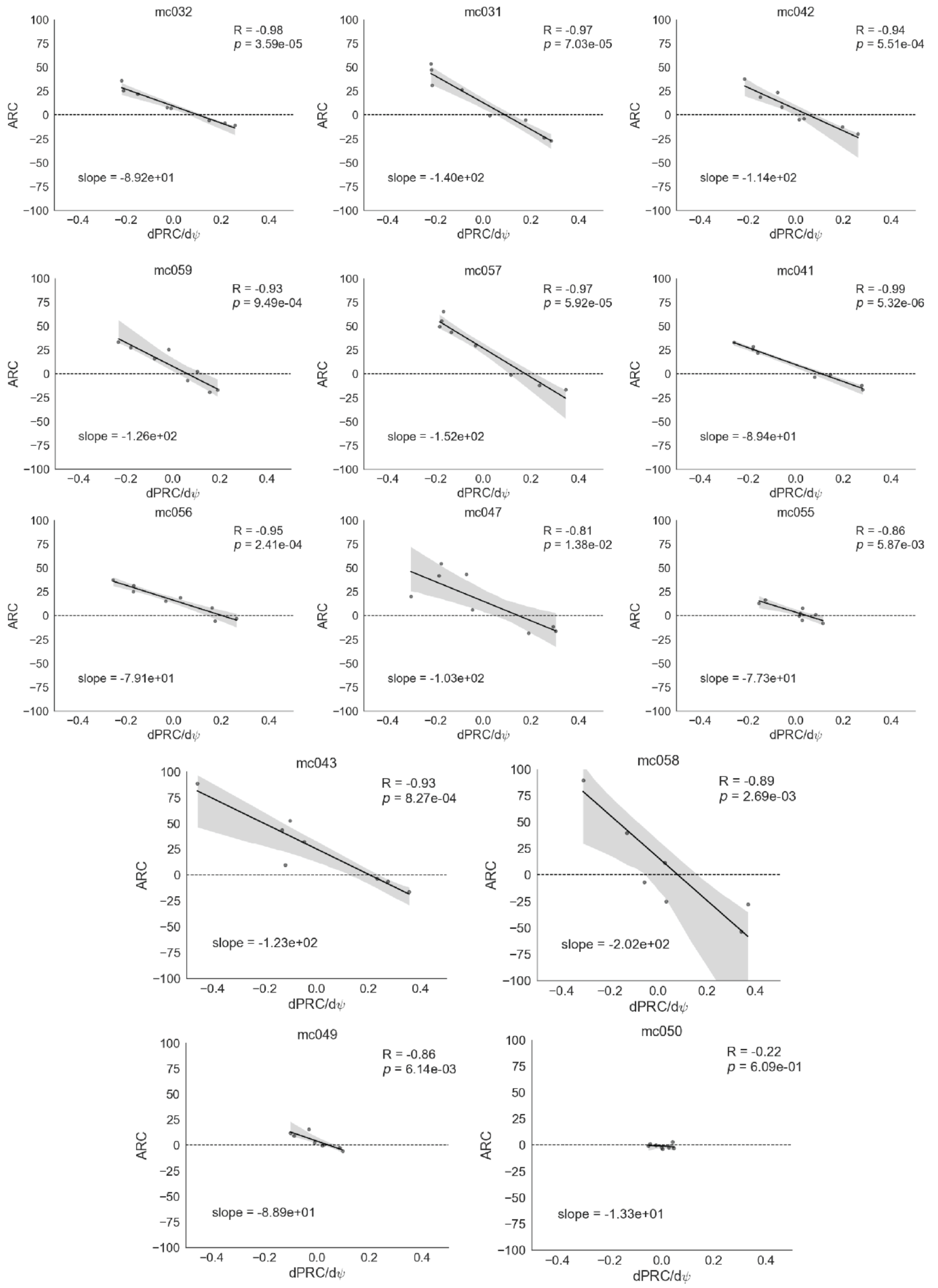


**Fig S2. Correlation between the ARC and the derivative of the PRC for individual animals.** Each plot displays the Pearson's correlation coefficient, the corresponding p-value from the statistical test, and the slope of the regression line.

### 2. Network tendencies in response to external perturbations

Another way of understanding how the instantaneous effect of stimulation translates into real changes in network synchrony is to consider the network's characteristic curve, which illustrates how steady state synchrony $\rho = \rho_{st}$ varies with intrinsic forces (Fig. S3, top). For illustration purposes, this curve was generated by varying the coupling $K$ (vertical axis) under no noise and a fixed $\gamma = 1$. However, similar curves, merely vertically shifted, can be generated for different noise levels and distribution widths. Accordingly, the vertical axis can be interpreted as a more general effective coupling that represents the balance of synchrony-promoting versus desynchronizing parameters. This characteristic curve defines the network's tendency to change its state in response to disturbances. In this context, stimulation acts as an external disturbance, the size of which is determined by the stimulation term in the reduced model. At low oscillation amplitudes, stimulation induces a large disturbance in coupled oscillators (indicated by arrows on the y-axis), but this disturbance translates into small changes in network synchrony (represented by the width of the red or green shades) due to the system's tendency in maintaining its asynchronous state. In addition, when the system is close to its lower bound, the same disturbance size leads to greater amplification compared to suppression (wider red shade than green). At higher amplitudes, small disturbance in oscillators and the network's characteristic curve both contribute to even lower changes in the state of synchrony. It is at intermediate values of oscillation amplitude that both the disturbance size and networks susceptibility to change are high, leading to more substantial changes in synchrony (the widest red and green shades).

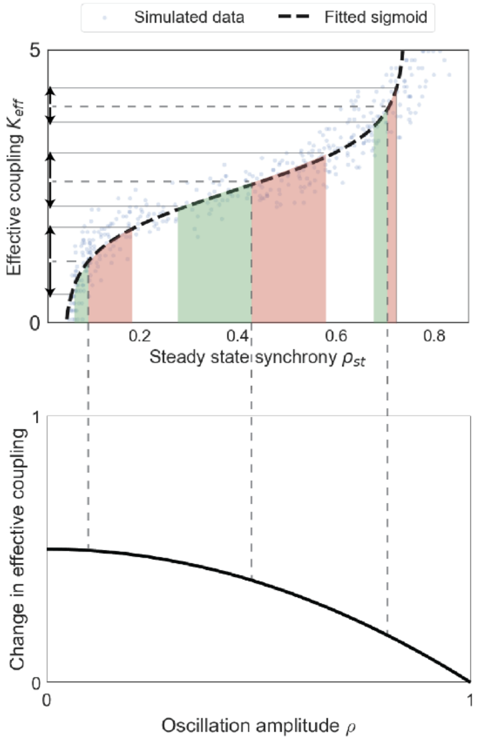


**Fig. S3. Role of network dynamics in the amplitude-dependence of the response** illustrated by a typical operating curve of a Kuramoto network (top) and stimulation-induced disturbance as a function of oscillation amplitude (bottom). The extent of change in network synchrony as a result of stimulation depends on the size of the induced disturbance as well as network tendencies at that specific amplitude.

#### 3. Model fitting results

The mean synchrony values arising from subject-specific best-fit models were presented in the main text. While those values exhibited relatively small variations, the exact values for the parameters of the underlying Kuramoto models varied substantially. This discrepancy likely stems from the limitations of parameter recovery in this mathematical model. To asses this, synthetic data generated by model simulations were fed into the fitting algorithm to determine whether the underlying parameters could be recovered. In a set of model fitting trials performed with synthetic data, we observed that the exact values of network parameters $K$, $\gamma$, and $\sigma$ cannot be recovered reliably. However, as shown in the main text, the mean synchrony can be consistently recovered.

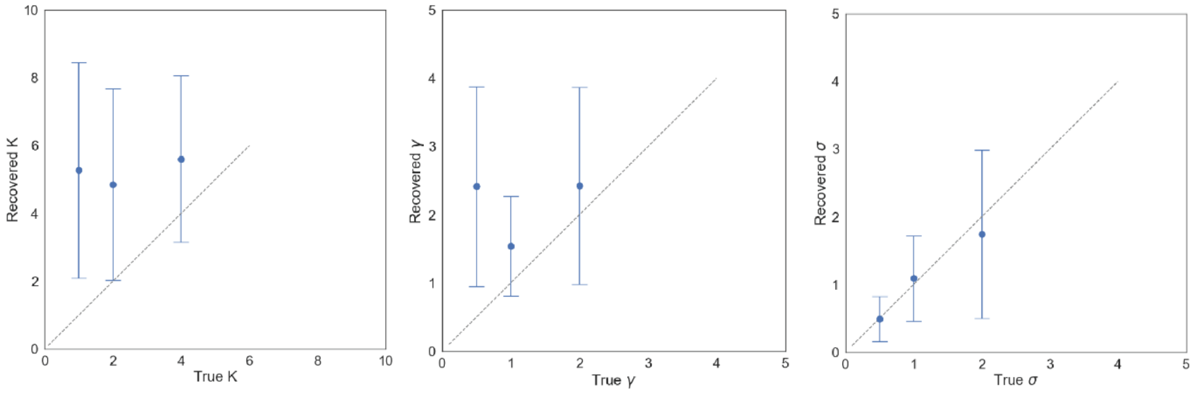


**Fig S4. Parameter recovery of the Kuramoto model using synthetic data.** A comparison of the true and recovered values for the intrinsic network parameters $K$, $\gamma$, and $\sigma$ reveals suboptimal recovery of these parameters.

One metric to quantify this feature of the model is the deviation from the critical coupling in the system. At the critical coupling, the network undergoes a transition from asynchronous state to partial synchrony and macroscopic clusters of entrained oscillators begin to emerge. It has been demonstrated that for a noise-free network with a Cauchy distribution of natural frequencies, the critical coupling is [1]:

$$K_c = 2\gamma.$$

For noisy network of coupled oscillators with the noise intensity of $D = \sigma^2/2$, this relationship takes the following form [2]:

$$K_c = 2\gamma + 2D.$$

As a result, $2\gamma + 2D - K$ can be viewed as a measure of deviation from the critical point, providing an indirect estimate of the system's synchrony level. Table S1 summarizes the results of the model fitting for all subjects, each with a set of three trials. As it can be seen from the table, the exact values of the mentioned model parameters vary even within animals. Nonetheless, the deviation from the critical point remained consistent across trials, which is further quantified by the coefficient of variation (CV) for this metric. It should be also noted that the goodness of the fits is presented in the table using the value of the minimized objective function $f$ which can be interpreted as $f = 1 - R^2$.

**Table S1. Results of the model fitting for individual animals.** The parameter values corresponding to the best fit at each optimization run are presented for three trials per animal.

| Subject | $K$ | $\gamma$ | $\sigma$ | $\omega_0$ | $f$ | Deviation | CV |
|---|---|---|---|---|---|---|---|
| mc031 | 5.6228 | 9.1736 | 3.7082 | 224.387 | 0.0331 | 26.47515 | |
| mc031 | 5.0726 | 9.9664 | 3.1844 | 223.9469 | 0.0357 | 25.0006 | |
| mc031 | 5.1193 | 9.4422 | 3.5884 | 222.7461 | 0.0356 | 26.64171 | 0.028323 |
| mc032 | 5.7694 | 0.2256 | 5.786 | 225.6911 | 0.0351 | 28.1596 | |
| mc032 | 9.4375 | 6.4934 | 4.8966 | 225.204 | 0.0345 | 27.52599 | |
| mc032 | 4.286 | 0.6135 | 5.639 | 225.5072 | 0.0378 | 28.73932 | 0.017607 |
| mc041 | 4.8273 | 10 | 3.4075 | 223.4505 | 0.0274 | 26.78376 | |
| mc041 | 2.6199 | 9.6109 | 3.1597 | 221.3587 | 0.0251 | 26.5856 | |
| mc041 | 0.7454 | 9.998 | 2.6151 | 222.1537 | 0.0267 | 26.08935 | 0.011027 |
| mc042 | 4.579 | 5.0208 | 4.7925 | 224.1004 | 0.0271 | 28.43066 | |
| mc042 | 5.1173 | 0 | 5.8324 | 222.9416 | 0.0284 | 28.89959 | |
| mc042 | 4.5156 | 9.9736 | 3.7427 | 225.5895 | 0.029 | 29.4394 | 0.01425 |
| mc043 | 5.5455 | 9.9726 | 3.9521 | 218.0733 | 0.0341 | 30.01879 | |
| mc043 | 6.1547 | 10 | 3.6659 | 218.4292 | 0.0336 | 27.28412 | |
| mc043 | 2.6475 | 0.8584 | 5.5632 | 218.2556 | 0.0375 | 30.01849 | 0.044287 |
| mc047 | 6.5101 | 1.5361 | 5.7613 | 215.0154 | 0.027 | 29.75468 | |
| mc047 | 8.0478 | 0.2487 | 5.8597 | 215.3317 | 0.0267 | 26.78568 | |
| mc047 | 9.7847 | 10 | 4.1124 | 215.8416 | 0.0296 | 27.12713 | 0.047562 |
| mc049 | 2.5799 | 9.9541 | 3.1715 | 221.1876 | 0.0324 | 27.38671 | |
| mc049 | 5.2138 | 9.3716 | 3.9774 | 220.747 | 0.0345 | 29.34911 | |
| mc049 | 0.0628 | 9.0564 | 3.4165 | 219.7357 | 0.0357 | 29.72247 | 0.035549 |
| mc050 | 9.5106 | 5.4175 | 5.2621 | 218.5247 | 0.0284 | 29.0141 | |
| mc050 | 0.7945 | 6.4858 | 3.4479 | 217.5699 | 0.0285 | 24.06511 | |
| mc050 | 9.3705 | 4.6673 | 5.1967 | 218.3789 | 0.029 | 26.96979 | 0.0761 |
| mc055 | 0.915 | 9.6541 | 3.2064 | 249.9889 | 0.0342 | 28.6742 | |
| mc055 | 2.4475 | 9.2304 | 3.6434 | 250 | 0.0325 | 29.28766 | |
| mc055 | 0.0054 | 6.7924 | 3.9697 | 249.0729 | 0.0349 | 29.33792 | 0.010369 |
| mc056 | 7.8124 | 4.3539 | 5.3867 | 248.5024 | 0.0329 | 29.91194 | |
| mc056 | 8.3298 | 10 | 4.1165 | 250 | 0.0325 | 28.61577 | |
| mc056 | 2.4395 | 8.6916 | 4.0708 | 250 | 0.0349 | 31.51511 | 0.03951 |
| mc057 | 1.3925 | 9.9441 | 1.4411 | 217.2204 | 0.0393 | 20.57247 | |
| mc057 | 6.2647 | 1.148 | 5.3233 | 219.1022 | 0.0431 | 24.36882 | |
| mc057 | 1.3622 | 9.9063 | 1.9662 | 219.1962 | 0.0409 | 22.31634 | 0.069207 |
| mc058 | 2.3443 | 6.0666 | 3.564 | 217.7543 | 0.0431 | 22.491 | |
| mc058 | 0 | 8.0559 | 2.4053 | 216.5805 | 0.0435 | 21.89727 | |
| mc058 | 0.0006 | 9.9595 | 0.7317 | 217.5308 | 0.0439 | 20.45372 | 0.039581 |
| mc059 | 8.511 | 10 | 4.4737 | 250 | 0.0453 | 31.50299 | |
| mc059 | 1.1251 | 5.7165 | 4.1351 | 249.9951 | 0.0454 | 27.40695 | |
| mc059 | 7.502 | 2.6516 | 5.4681 | 250 | 0.0433 | 27.70132 | 0.064612 |

### 4. Analytical estimation of the average network synchrony

It has been demonstrated that assuming a linear relationship between the abstract network activity and the experimentally measured signal, Hilbert transform of the signal can be used to relate the experimental data to the model output by:

$$a = C.\rho \,, \psi_e = \psi, \tag{1}$$

where $a$ and $\psi_e$ are the envelope amplitude and instantaneous phase derived from the Hilbert transform of the signal, respectively, and $C$ represents a constant relating experimental amplitude to network synchrony. Given this relationship, the oscillation phase and amplitude embedded in the order parameter were used as the basis for predictions. In order to translate the model's predictions, an estimate for $C$ is required to convert the signal values into a range of synchrony.

It has been shown that when $Z(\theta)$ contains only the q$^{th}$ harmonic, the instantaneous ARC and PRC (introduced in Section 2.3 of the main text as $\mathbf{P}(\psi) = \frac{I}{2}(1-\rho^2)\cos(\psi)$ and $\mathbf{\Psi} = \frac{I}{2\rho}(1+\rho^2)\sin(\psi)$) averaged across all amplitudes are correlated as follows [3]:

$$\overline{\mathbf{P}}\,(\psi) = -\frac{\nu_q}{q\tilde{\nu}_q}\frac{d\overline{\mathbf{\Psi}}}{d\psi} \;\Rightarrow \overline{\mathbf{P}}\,(\psi) = -\alpha_{th}\frac{d\overline{\mathbf{\Psi}}}{d\psi}\,, \tag{2}$$

where

$$\nu_q = \int_0^1 h(\rho)(1-\rho^2)\rho^{q-1}d\rho\,, \tag{3}$$

$$\tilde{\nu}_q = \int_0^1 h(\rho)(1+\rho^{-2})\rho^{q}d\rho\,. \tag{4}$$

Here, $h(\rho)$ represents the probability density function (PDF) for the system having an amplitude of $\rho$. In the case of dominant first harmonic, $q = 1$, which aligns with the model's assumption and approximates many experimentally reported response curves for neurons [4,5], the relationship slope $\alpha_{th}$ can be further simplified to:

$$\alpha_{th} = \frac{\int_0^1 h(\rho)(1-\rho^2)d\rho}{\int_0^1 h(\rho)(\rho+\rho^{-1})d\rho}\,. \tag{5}$$

On the other hand, the observed correlation between the block-based ARC and the derivative of the block-based PRC (Fig. 4 in main text) allows us to write:

$$ARC_b = -\alpha_{exp}\,\frac{d(PRC_b)}{d\psi}. \tag{6}$$

As mentioned above, due to the different scales of network synchrony and the experimentally measured signal, the experimental slope is also proportionally scaled:

$$a = \frac{\rho}{C} \Rightarrow ARC_b \approx \overline{\boldsymbol{P}}\,(\psi)/C, \tag{7}$$

Therefore, predicting the experimental slope requires knowing the constant of proportionality $C$, which yields the following equation for the experimental slope:

$$\alpha_{exp} = \frac{1}{C}\cdot\frac{\int_0^{1/C} h(Ca)(1-C^2a^2)Cda}{\int_0^{1/C} h(Ca)(1+C^{-2}a^{-2})CaCda}. \tag{8}$$

The above equation describes how the experimental slope varies as a function of $C$ given a specific $h(a)$ which can be extracted from the animal's ECoG recordings. For each subject, this equation was solved numerically in MATLAB using the corresponding amplitude PDF derived from off-stimulation recordings. Fig. S5 illustrates how mean synchrony is expected to vary as a function of the slope under various amplitude PDFs for different animals. The empirically determined slope value is indicated by a vertical grey ribbon in each plot. Overall, the results indicated that these slope values suggest underlying networks with low sustained synchrony. However, due to the assumptions inherent in the analytical derivations and the deviations of the experimental curves from the analytical ones due to their definitions, we proceeded with model fitting as the primary method for determining network synchronies.

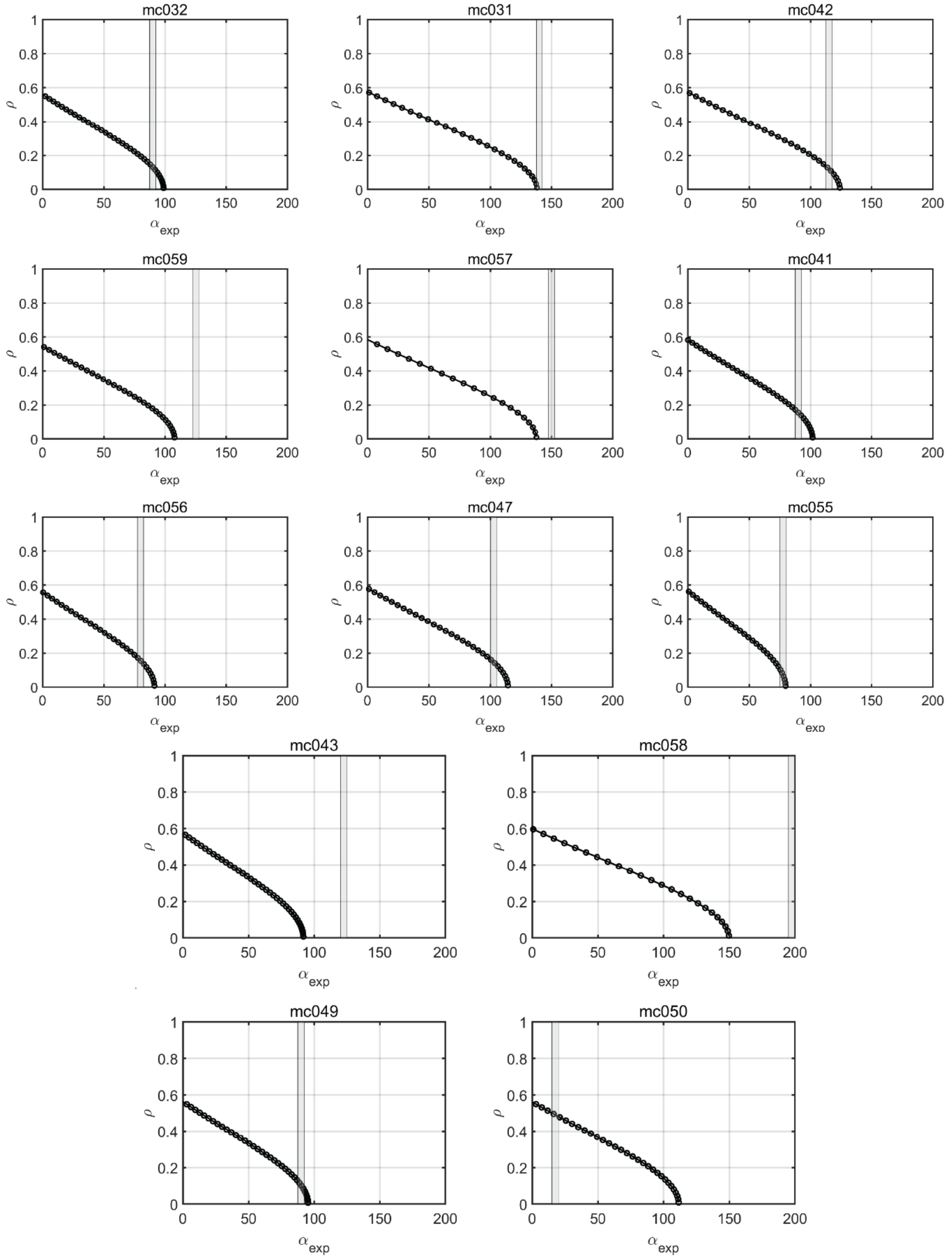


**Fig S5. Analytically derived variation of mean network synchrony as a function of the correlation slope under different amplitude PDFs.** For each animal, the empirically determined slope value is indicated by a thin vertical ribbon. Most of these empirical values fall within the region that implies underlying networks with low synchrony.

In order to develop a quick first-order analytical estimate for this slope, one can assume that the system dynamics are summarized by an average synchrony, meaning that the whole probability function is concentrated at the mean value $h(\rho) \approx \delta(\bar{\rho})$. Under this condition,

$$\alpha_{th} = \frac{1-\bar{\rho}^2}{\bar{\rho}+\bar{\rho}^{-1}}\,, \tag{9}$$

which implies that $\alpha_{th}$ lies in the range of [0 0.3]. Using the proportionality constant between the model's synchrony $\rho$ and signal's envelope amplitude $a$, the relationship between the experimental slope and network synchrony can be derived as follows:

$$\alpha_{exp} = \frac{1}{C} \cdot \frac{\bar{\rho}-\bar{\rho}^3}{\bar{\rho}^2+1}\,. \tag{10}$$

It is insightful to examine the above relationship at the extremes of low and high synchrony to get a better understanding of the variations of the experimental slope:

$$\begin{cases} \bar{\rho} \to 0 \;\Rightarrow\; \alpha_{exp} \approx \bar{a} \\ \bar{\rho} \to 1 \;\Rightarrow\; \alpha_{exp} \approx 0 \end{cases} \tag{11}$$

This reveals an interesting trend implying that as the network exhibits lower levels of sustained synchrony, the experimental slope approaches the value of the average amplitude. Additionally, these estimates are independent of $C$ and they eliminate the need for model fitting to extract the mean synchrony if only the limits are of interest. Nonetheless, the full analytical relationship given by eq. (11) still relies on the proportionality constant between the experimental data and model's synchrony to relate the corresponding slopes. To circumvent this challenge, normalized ARCs can be employed where both response curves demonstrate change in amplitude divided by the amplitude itself.

$$\frac{\bar{\mathbf{P}}(\psi)}{\rho} = -\frac{(1-\bar{\rho}^2)}{(1+\bar{\rho}^{-2})\bar{\rho}^2}\frac{d\bar{\mathbf{\Psi}}}{d\psi} \;\Rightarrow \tilde{\alpha}_{th} = \frac{1-\bar{\rho}^2}{\bar{\rho}^2+1}, \tag{12}$$

$$\frac{ARC_b}{a} = -\tilde{\alpha}_{exp}\,\frac{d(PRC_b)}{d\psi}. \tag{13}$$

This leads to comparable normalized slopes:

$$\frac{\bar{\mathbf{P}}(\psi)}{\rho} = \frac{ARC_b/C}{a/C} = \frac{ARC_b}{a} \Rightarrow \tilde{\alpha}_{th} = \tilde{\alpha}_{exp}. \tag{14}$$

Again, looking at the limits offers an insight into the behaviour of the experimental slope as a function of network synchrony.

$$\begin{cases} \bar{\rho} \to 0 \;\Rightarrow\; \tilde{\alpha}_{exp} \approx 1 \\ \bar{\rho} \to 1 \;\Rightarrow\; \tilde{\alpha}_{exp} \approx 0 \end{cases} \tag{15}$$

This characteristic behaviour of the slope provides a generalized analytical metric to develop a sense about the network synchrony by looking at the experimental correlation between normalized ARC and derivative of the PRC. Fig. S6 summarizes the value of this normalized slope for different subjects.

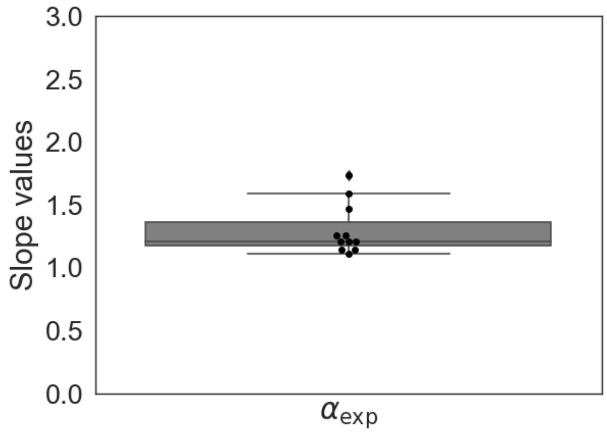


**Fig S6. Slope values for the relationship between ARCs and PRC derivatives in different subjects.** Values close to 1 indicate that the underlying networks feature very low sustained synchrony.

Again, while most values suggest low-synchrony networks by being close to 1, their exact values lack sufficient reliability to serve as a basis for further analyses. This is mainly due to the crude way of defining experimental ARC and PRC

.

### 5. Baseline subtracted suppression and amplification profiles

Given the fact that noisy systems with fluctuating states (such as a network with varying synchrony) tend to regress to the mean, it's important to account for this phenomenon when assessing changes due to stimulation. The reference lines shown in the main text illustrate this regression tendency in the absence of stimulation. By calculating the difference between the suppression/amplification line and the reference line, one can determine the net effects of stimulation. For the block-based approach, the reference trend line was subtracted from the amplification/suppression datapoints. The regression lines for reference-subtracted values, depicted in Fig. S7A indicated significant increase of the effect (slope=-0.152, *p*-value=1.7e-6) in suppression versus nonsignificant change in amplification (slope=0.067, *p*-value=0.49). Also, Fig. S7B shows trends for the pulse-based method where the reference curve is subtracted from the original amplification/suppression, reflecting a general decay in the effect at very low synchrony values.

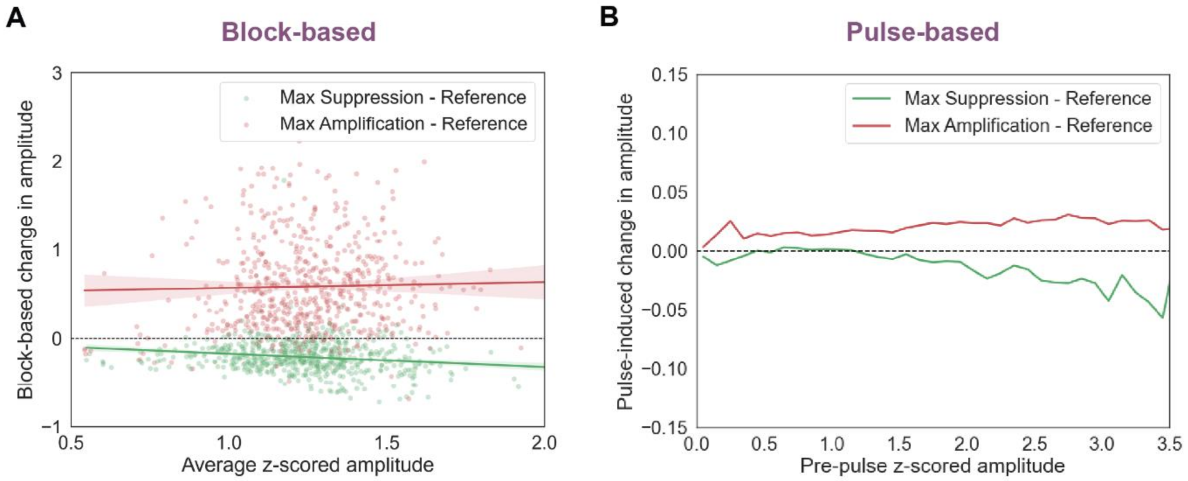


**Fig S7. Reference-subtracted trends for amplification and suppression.** Both block-based (A) and pulse-based (B) approaches capture a diminished stimulation effect.
